# Robust nomenclature and software for enhanced reproducibility in molecular modeling of small molecules

**DOI:** 10.1101/429530

**Authors:** Hesam Dashti, Jonathan R. Wedell, Gabriel Cornilescu, Charles D. Schwieters, William M. Westler, John L. Markley, Hamid R. Eghbalnia

## Abstract

Computational molecular dynamics, energy minimization, and modeling of molecular interactions are widely used in studies involving natural products, metabolites, and drugs. Manually directed computational steps commonly utilize an evolving collection of experimental and computational data, to which new data sources are added or modified as needed. Several software packages capable of incorporating sources of data are available, but the process remains error prone owing to the complexities of preparing and maintaining a consistent set of input files and the proper post-processing of derived data. We have devised a methodology and implemented it using an extensible software pipeline called RUNER (for Robust and Unique Nomenclature for Enhanced Reproducibility) that creates a robust and standardized computational process. The pipeline combines a web service and a graphical user interface (GUI) to enable seamless modifications and verified maintenance of atom force field parameters. The GUI provides an implementation for the widely used molecular modeling software package Xplor-NIH. We describe the RUNER software and demonstrate the rationale for the pipeline through examples of structural studies of small molecules and natural products. The software, pipeline, force field parameters, and file verification data for more than 4,100 compounds (including FDA-approved drugs and natural products) are freely accessible from [http://runer.nmrfam.wisc.edu].

**Author Summary:** We describe an automated and verifiable computational pipeline for calculating the force field parameters of small molecules. The pipeline integrates several software tools and guarantees reproducibility of the parameters by utilizing a standard nomenclature across multiple computational steps and by maintaining file verification identifiers. We demonstrate the application of this pipeline to (a) processing of more than 4,100 compounds in high-throughput mode, and (b) structural studies of natural products. The graphical user interface (GUI) associated with the pipeline facilitates the manually tedious steps of force field parameters adjustments and supports visualization of the process.

## Introduction

Experimental data and computational modeling are complementary and often used together in a variety of studies. Experimental methods, such as X-ray crystallography and NMR spectroscopy, play key roles in determining structures, dynamic properties, and molecular interactions. Quantum chemistry and computational analyses are used to probe the energy landscapes of molecules, to calculate or refine their structures, and to model their dynamic and spectroscopic parameters. For example, atom-specific experimental data (e.g., NMR residual dipolar couplings, nuclear Overhauser effect restraints, three-dimensional coordinates derived from X-ray density)^1,2^ are used as input to molecular modeling software suites such as Xplor-NIH.^3,4^ Such data can form the basis for determining structures of small molecules and for studying their interactions with macromolecules. The diversity of input data types and objectives of the studies yield to a variety of starting points for studies. The structure of the target macromolecule – a macromolecule with an entry in the Protein Data Bank (PDB), or a protein under investigation – can provide a starting point for molecular modeling. At the same time, one or more small molecules can be selected as bait for exploring potential interactions. One way to assess macromolecule-ligand interactions is through computer simulations using molecular docking software that scores interactions.^5^ More commonplace is the use of a combination of both computational and experimental approaches in investigations involving molecular fragments or other small organic molecules (commonly known as metabolites or ligands).^4,6-10^ Several existing software packages facilitate docking studies of ligands and macromolecules (for examples, see^11,12^). If the goal is to identify which molecules from a set of compounds are likely to bind to a given macromolecule, interaction scores are determined, and those ligands with the highest scores are selected for further experimental investigation. The approach is iterative, and the workflow remains fluid in order to accommodate modifications in the use of computational or experimental evidence as the data accumulate (Figure 1).

**Figure 1.**
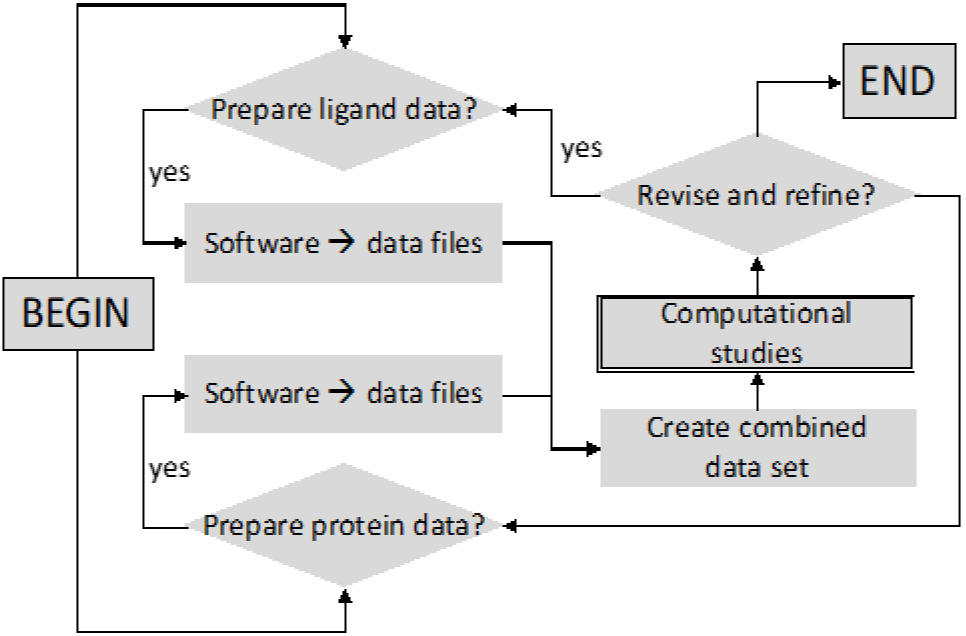
Iterative workflow of molecular modeling. In an iterative modeling approach, the same logical steps may be repeated by using different software, or different versions of the same software, with or without additional data. A protocol to verify the consistency and reproducibility of the results must incorporate reproducible and verifiable data and processes.

Such calculations can be laborious and highly error-prone, because they typically involve multiple software tools with differing input-output requirements, which are “pipelined” such that the output of one tool is fed as the input to the next tool in the workflow. What is needed is a protocol that guarantees consistency across the iterative deployment of multiple software tools having different input-output requirements, restraint settings, and other computational factors.

Consistency of nomenclature is vital to scientific communication. In a modular pipeline-based approach, communication through properly formatted input in each step of data processing is critical to the integrity of results. In the pipeline, the improper output of a single step can render the pipeline irreproducible. Therefore, communication of nomenclature and format must be carefully crafted and controlled across multiple iterative steps in biochemical and biomedical studies. Although atom naming in proteins and nucleic acids has been standardized,^13^ one of the main challenges in the utilization of integrative/hybrid methods is the lack of consistent atom labels for small molecules. If different experimental and computational tools utilize different atom nomenclature, the task of converting the output and atom labels from one step to the input labels for the next step is tedious and error-prone.^14^

Moreover, while consistent nomenclature is necessary, it is not sufficient. Whereas the atomic force field parameters for amino acids and nucleic acids in macromolecules are rigorously calculated and predefined, the conformational diversity and the massive number of configurations of small molecules prohibits pre-calculation of these parameters. In fact, the possible combinatorial structures of synthesized or naturally produced small molecules is yet to be determined.^15,16^ Thus, the lack of standardized calculated force fields for small molecules must also be addressed.

Rigorous quantum mechanical calculations of parameters are computationally expensive; therefore, a number of software packages provide force field parameters derived from semi-empirical quantum mechanical computations as a means of accelerating the process.^17-22^ Text-processing wrappers have been developed, that translate the nomenclature of the modeled force field output to the nomenclature and data format used by different molecular modeling software packages.^23^ These wrappers are helpful, but they have technical and computational shortcomings. On the technical side, the integration of iterative steps is complicated because the wrappers do not use unique and reproducible atom nomenclature; moreover, not all wrappers are provided in the public domain. On the computational side, the use of different software packages in the workflow may require the transformation of data from multiple formats to multiple formats, a formidable data management problem. These problems can be compounded when atom nomenclature and/or data formats change from one software release to the next. Users may also initiate model changes that alter the data content. In summary, several factors can compromise the reproducibility of computations: a) differences in stereochemistry, b) differences in nomenclature, c) incompatibility between wrappers, d) user-initiated model changes that are not captured in an iterative workflow, and d) changes introduced across software releases.

The RUNER methodology is designed to address these challenges. To ensure the uniqueness and reproducibility of all atom names in experimental and computational data sets, we utilize ALATIS,^24^ a software package that takes a structure file as input and extends the system of international chemical identifiers^25^ (InChI) to enable the assignment of unique labels to all atoms. This step guarantees that all subsequent atom label transformations in the pipeline can be automatically managed without the need to perform many-to-many data transformations (Figure 2).

**Figure 2.**
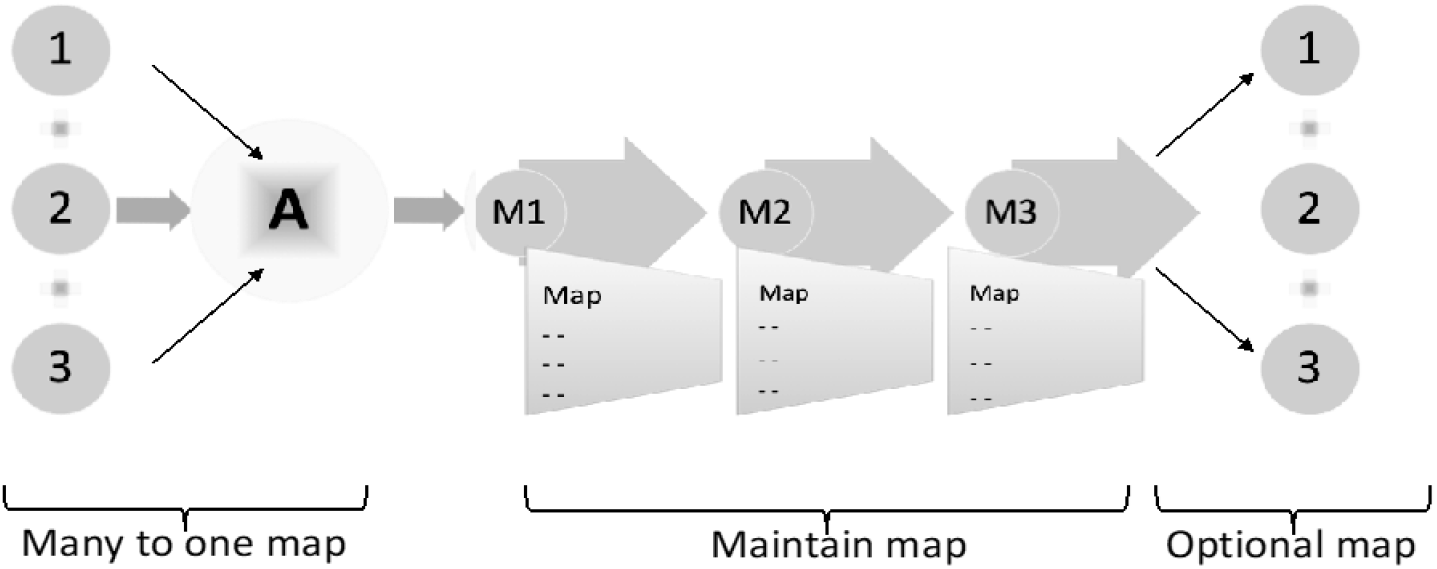
Managing data transformations. The RUNER workflow creates a map based on the ALATIS nomenclature (shown as A in the figure). A map is generated for each software package or data file as needed (shown as 1, 2, 3 in the left side of the figure). The map is maintained throughout the process (shown as M1, M2, M3), and the results are transformed back to the original nomenclature if necessary. In addition, in order to verify the integrity of the generated maps and data files, the process creates and maintains verification keys that are validated by software utilities in the computational pipeline.

## Methods

The algorithm that manages label transformations uses an internally maintained map between software-specific labeling and ALATIS nomenclature. All intermediate files maintain MD5 hash verification keys recorded in NMR-STAR format.^26^ RUNER validates the keys and generates a report, which warns users of any potential inconsistencies. Users can obtain results in either software-specific or ALATIS nomenclature, and they can obtain the map that controls the transformation to guarantee future reproducibility. The pipeline supports the commonly used molecular mechanics accessory package antechamber,^17,18^ an Amber tool^27^, which generates input files according to the general Amber force fields (GAFF and GAFF2) for the commonly used molecular modeling software suites CHARMM^7,8^ and Amber.^27,28^ As an alternative to the semi-empirical quantum chemistry (sqm) in antechamber, the molecular orbital package (MOPAC)^21,22^ is incorporated to the RUNER pipeline for faster atom charge calculations. The academic release of MOPAC software suite utilizes the favorably reviewed PM7 method for calculating atom charges that is used in the RUNER pipeline.^29,30^ We have developed a web server [http://runer.nmrfam.wisc.edu/] that provides free and unlimited access to this module of RUNER’s pipeline.

For the case of Xplor-NIH,^3,4^ which is widely used for structure calculations and investigations of ligand-binding of drug-like compounds, RUNER pipeline is equipped with text-processing modules that convert Amber force fields file formats to Xplor-NIH file formats. In addition, RUNER deploys a standalone graphical user interface (GUI) that enables the seamless inspection and modification of force field topology and parameters. We describe the different modules of the RUNER pipeline below in the “Design and implementation” section. Although the software components deployed in the current implementation of the RUNER pipeline represent archetypal tools commonly used in the field, RUNER is not limited to this specific set of software. RUNER can be adapted for use with other software components. In the “Results” section, we illustrate two applications of the methodology implemented by the RUNER pipeline.

### Design and implementation

RUNER (Figure 3) serves as computational pipeline that manages the consistency of model definition across computational molecular modeling stages. The system is composed of a web server module and a desktop standalone GUI. The web service generates formatted topology files, parameter files, and mapping data. Mapping data provide the basis for creating internal tables which guarantee the correct transformation of nomenclature among all experimental (e.g. structural restraints derived from NMR experiments) and computational data sources. Additionally, the mapping data can be used to convert the nomenclature in the format desired by the user.

**Figure 3.**
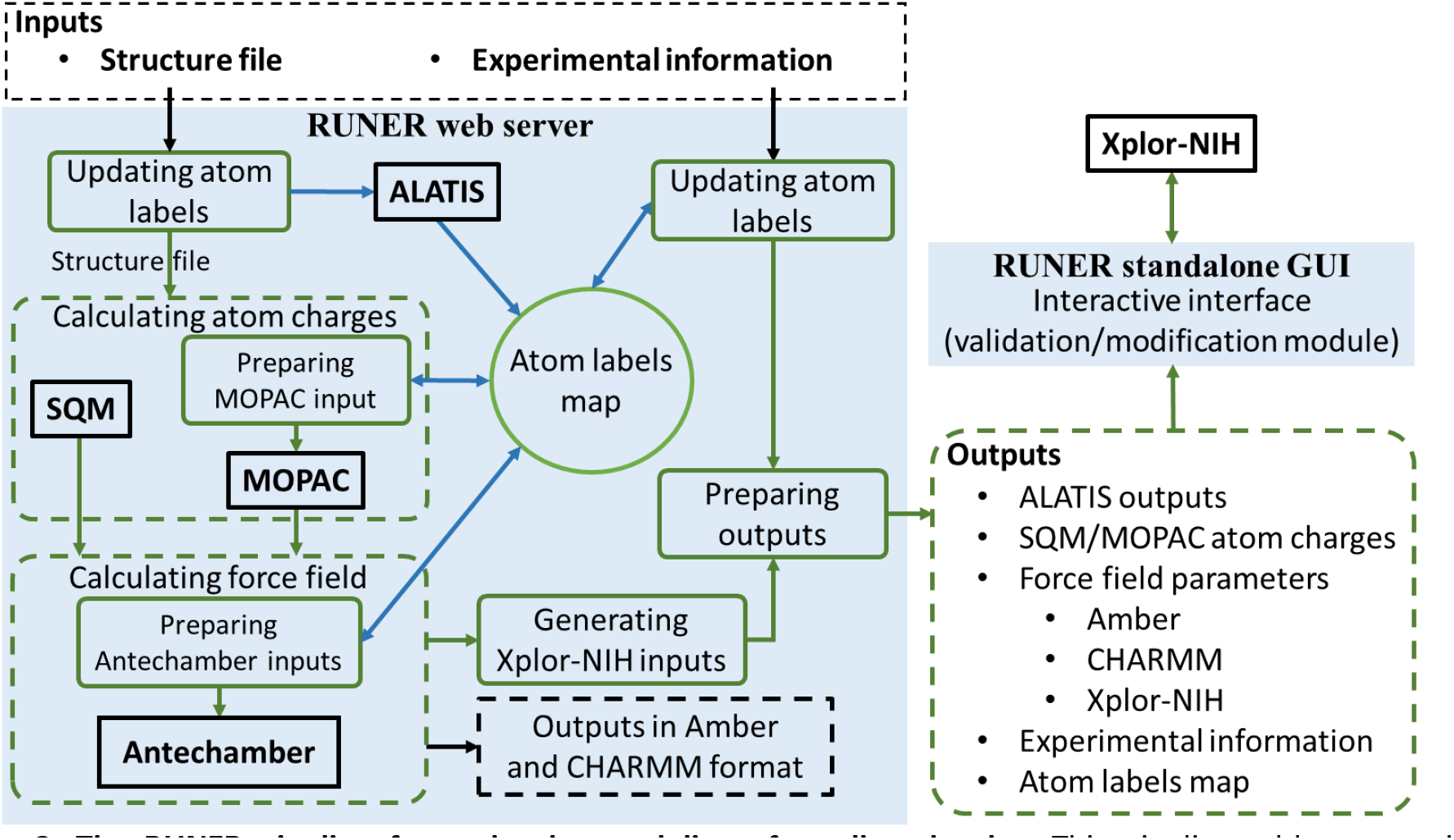
The RUNER pipeline for molecular modeling of small molecules. This pipeline addresses major challenges involved in the utilization of hybrid methods in computational studies of small molecules. Text processing modules of the pipeline are shown in green boxes. Green arrows indicate directions of workflows in the RUNER pipeline. Blue lines show construction and retrieval of the atom labels map. The dashed-lines indicate groups of entities (computational modules or files), and black boxes indicate external software programs. The service and the software are available at [http://runer.nmrfam.wisc.edu/].

Currently, two software packages (antechamber and MOPAC) are utilized to generate the required force field parameters. Although these software packages are sufficient for the majority of small molecules and natural products, other software programs can be incorporated in the same framework. The outputs of the web server can be used as input to several software packages, including Xplor-NIH, CHARMM, and Amber. The desktop GUI streamlines the application of hybrid computational molecular modeling within the Xplor-NIH environment.

### Main features and key algorithms of the RUNER web server

• **Inputs**. The web server module of the pipeline implements a series of input structure files. This module utilizes input structure files in MOL V2000 or its corresponding Simple Data Format (SDF).^31^ ChemDraw Exchange (CDX) and Protein Data Bank (PDB) formats are also accepted by the web server, which utilizes Open Babel^32^ to interpret and convert these formats. The web server utilizes Open Babel tools to generate three-dimensional coordinates from two-dimensional structures and to add explicit hydrogen atoms when needed. In addition to the structure file, the web server accepts data from NMR experiments, including residual dipolar coupling (RDC), nuclear Overhauser effect (NOE), and dihedral angle restraints.

- **Updating atom labels**. The ALATIS software in this module is used to generate unique atom labels based on the input molecular structure files, which are then used to update the atom labels in the uploaded experimental restraints and to generate a new structure file according to the unique atoms.
- **Atom labels map**. Simple data formats for storing molecular structures (like MOL and SDF) do not allow assigning customized atom labels, and atoms in these formats are referred to according to the sequence in which the atoms are defined in the file. For example, the first defined atom is labeled 1, the next defined atom is labeled 2, and so on. To accommodate these widely used data formats for small molecules, ALATIS rearranges the atom definitions to assign unique labels to the atoms. Alternatively, in more enhanced data formats, such as the PDB format, a column with four digits is considered for customized atom names [PDB v3.3]. During calculations of the force field parameters, atoms need to be assigned to distinct labels. This is because, atoms will be clustered into groups, based on their adjacent atoms and bonds, and assigning different atoms to the same label confuses the clustering processes such that force field parameters cannot be provided for every individual atom. Taking these properties and constraints into account, the RUNER web server constructs a mapping from the input atom indices (MOL or SDF) and atom labels (PDB) to the unique atom labels generated by ALATIS. This map is utilized to modify input experimental information. When MOPAC is utilized in the pipeline, it is necessary to prepare an input file in MOPAC data format. To prepare the input file and also for interpreting MOPAC outputs, RUNER uses the atom labels map to preserve the consistency between the input and output files. Users who want to preserve the input atom nomenclature in the output can specify this before submitting a job to the RUNER web server. In such cases, the ALATIS naming system is used in the background in order to manage the data transformations, and the atom labels map is used to convert all output atom labels back to the input atom labels.
- **Calculating force field**. MOPAC outputs cannot be directly utilized to prepare force field parameters necessary in the widely used Xplor-NIH program. In RUNER’s web server MOPAC and antechamber are pipelined such that atom charge outputs of MOPAC are converted to standard antechamber input format. Antechamber then uses these charges (from SQM or MOPAC) to calculate force field parameters in Amber format.
- **Outputs of antechamber**. Antechamber generates atom force field parameters in Amber and CHARMM format, and these formats are included in the output of the web server. Because the input file to antechamber is a structure file with unique atom names, the force field output parameters of antechamber (in Amber and CHARMM formats) are defined in terms of these unique atom names.
- **Generating Xplor-NIH inputs**. Antechamber generates a parameter-topology specification file known as a “prmtop file” (see the online documentation at http://ambermd.org/prmtop.pdf and http://ambermd.org/formats.html). The RUNER web server is equipped with a text-proessing module (Figure 3) that processes the prmtop file and generates formatted topology and parameter files to be used by Xplor-NIH. While generating the topology and parameter files, RUNER resolves three demands of Xplor-NIH that are generally ignored by antechamber:

I. **Chiral proper dihedral angles:** Xplor-NIH requires the listing of chiral proper dihedral angles as improper dihedral angles. RUNER utilizes the standard InChI string generated for the compound by ALATIS to identify its chiral centers and applies the corresponding improper syntax. The representations of chiral centers in InChI strings are discussed in references.^24,25^
II. **Planar proper dihedral angles:** Xplor-NIH requires that the proper dihedral angles that enforce planarity are defined as improper dihedral angles, and the RUNER web server identifies them and fulfills this requirement.
III. **Replication of information:** Because the parameters corresponding to a particular topology are constant values, regardless of the way the set is defined (for example an angle can be defined as a sequence of atoms C1-C2-C3 or C3-C2-C1), Xplor-NIH requires a single definition for each topology. The RUNER web server finds and removes any duplicates.
- **Creation of validation records**. A 128 bit MD5 hash number is used to validate model files at any time.^33^ The validation record is maintained alongside the model files, and records are validated by using RUNER utilities.
- **Preparing outputs**. This module collects the results from different modules and prepare a compressed file to be displayed on the website.
- **Outputs**. The results on an output webpage are organized based on their application in Amber, CHARMM, Xplor-NIH software programs.

### Main features and key algorithms of the RUNER standalone GUI

Depending on the structural particularities of the compound of interest, one may need to use the validation/modification module in RUNER’s pipeline (Figure 3) to remove, add, or adjust various topology/parameter statements. Because the definition of atoms, bonds, bond angles, proper and improper dihedral angles, and non-bonded terms and their corresponding force field parameters are distributed in two files, it is difficult to make such adjustments. To facilitate these modifications, we have created a standalone desktop GUI as part of the RUNER pipeline (**Supporting materials**). The main window of the GUI contains three panels with tables to list the proper/improper dihedral angles, bonds, and bond angles and allows direct modification of the associated force field parameters. In addition, the GUI provides a module to investigate van der Waals statements and to modify Lennard-Jones parameters of non-bonded atoms with the same atom types, and also to create and adjust these parameters for non-bonded atoms with distinct atom types. The GUI allows immediate localization of atoms in the context of the three-dimensional structure of the molecule.

## Results

We demonstrate two applications of the RUNER computational pipeline: (1) the high-throughput processing of structures of compounds from databases (drugs and natural products) to obtain topology and parameter files needed for molecular interaction studies, and (2) streamlining the solution structure determination of natural products containing multiple chiral centers from NMR data by the use of a progressive stereo locking (PSL) strategy.^1^ The first application illustrates how RUNER can be used to automatically generate reproducible molecular information for a large data set, and the second application describes the needs and advantages of utilizations of the RUNER standalone GUI in structural studies of natural products.

### (1) Application of the RUNER pipeline to compounds from databases

To illustrate the high-throughput generation of reproducible atom force field parameters for compounds, we chose the set of FDA approved drugs from Selleckchem [http://www.selleckchem.com/] and the set of natural products from StreptomeDB 2.0.^34^ The structure files available from these two target databases were used as input to RUNER. Because many of the structure files archived in these databases only represent the heavy atoms of the compounds (with explicit hydrogen atoms missing), we utilized the Open Babel software package to preprocess the structure files to generate their complete 3D structures. Figure 4 shows a histogram of the number of atoms in each of the 3D structures of the 4,126 small molecules generated by Open Babel (1,342 drugs from Selleckchem and 2,784 natural products from StreptomeDB 2.0). The computational cycles required by the RUNER web server were provided by the high-throughput computing platform HTCondor,^35^ which supports multiple concurrent submissions. Using the HTCondor system enabled us to reduce the time for the entire calculations from approximately one year to about one day. The results are available on our website [http://runer.nmrfam.wisc.edu/] and are accessible through a search engine that is embedded in the RUNER website. On this website, for every compound, we provide a) the unique compound names and complete atoms labels (including hydrogens) as produced by ALATIS, b) a PDB file with three-dimensional coordinates for the uniquely-named atoms, c) input force fields files for molecular modeling with Amber, Xplor-NIH, and CHARMM, and d) explanatory meta-data regarding the compounds (e.g., targets of the drugs, formula, molecular weight, CAS Number, ALogP) as provided by the target databases.

**Figure 4.**
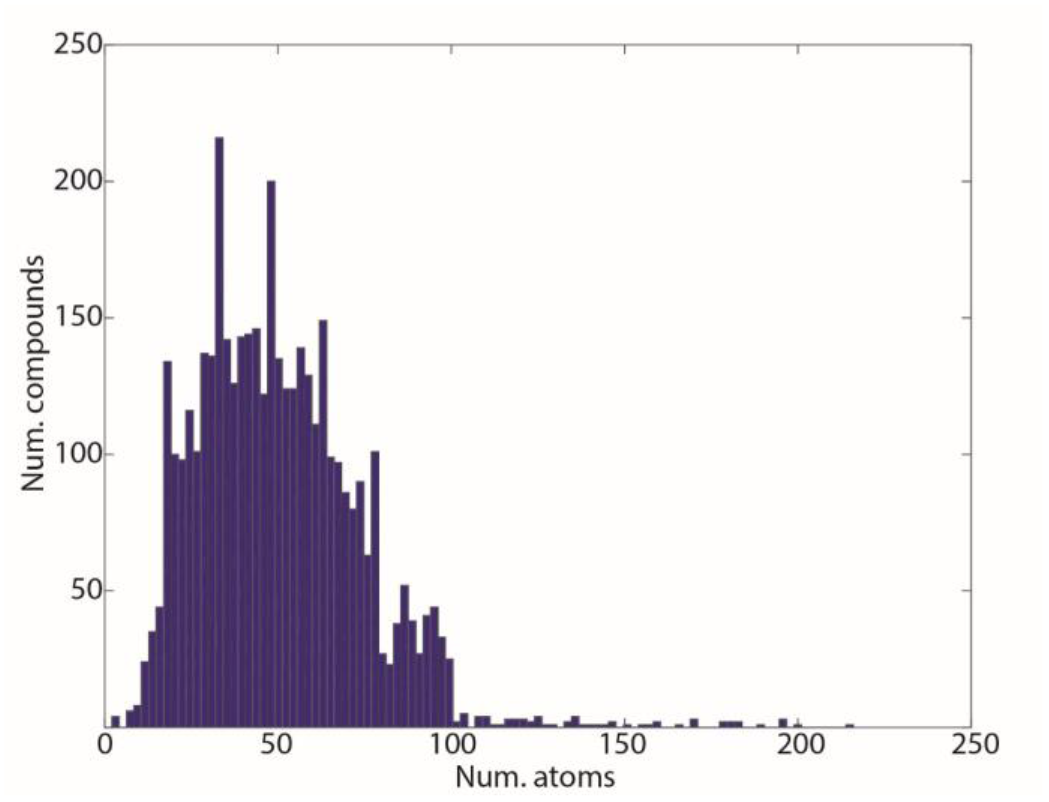
Histogram showing the number of atoms in the processed small molecules. The *x*-axis shows the number of atoms and the *y*-axis shows the corresponding number of compounds.

### (2) Application of the RUNER pipeline to natural product structure determinations

We selected two examples to illustrate how the RUNER pipeline was able to streamline the manual steps taken in determining configurations of natural products from NMR data by the Progressive Stereo Locking (PSL) method: 10-epi-8-deoxycumambrin B (38 atoms) and fibrosterol sulfate A (161 atoms). In addition, this example shows applications of the standalone GUI for seamless modifications of the force field parameters as needed for structural investigations.

**Deoxycumambrin**. NMR residual dipolar coupling restraints (^1^D_CH_ RDC) were utilized in determining the stereochemistry of this compound. Figure 5a shows the process of preparing input files to Xplor-NIH from the experimental restraints and force field parameters generated by PRODRG.^19^ Aggregation of the atom specific information from the NMR RDC restraints with the force field parameters, required keeping the atom labels consistent.

**Figure 5.**
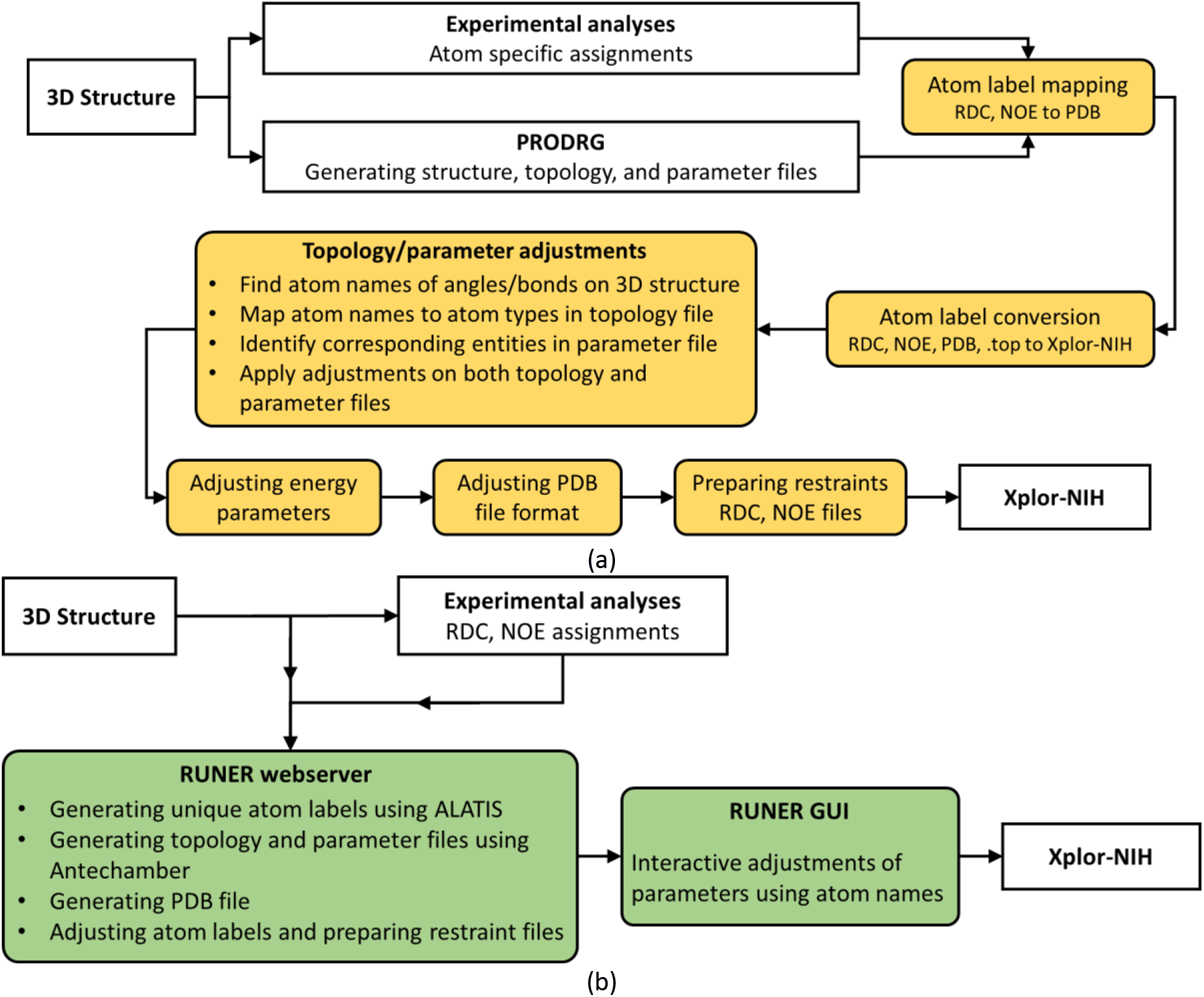
The process for structural investigation of natural products. (a) Conventional workflow as described in the PSL publication. In this workflow, the manual steps are shown in colored boxes. (b) Automation of the PSL steps in the RUNER pipeline. The green boxes show RUNER modules.

The ‘*Atom label mapping*’ step involved manually mapping the atom labels used in NMR RDC assignments (from the input 3D structure) to those provided by the PRODRG web server. In addition, because Xplor-NIH does not accept some atom labels generated by PRODRG, a second manual step (‘*Atom label conversion*’) was needed to convert the PRODRG atom labels to the standard accepted by Xplor-NIH. We note that, PRODRG only changes labels of hydrogen atoms, and the steps for mapping and converting atom labels needed to be performed manually on the 20 hydrogen atoms in the 10-epi-8-deoxycumambrin B molecule. Because force field parameters are stored in a specific data format in the topology (.top) and parameter (.par) files, every necessary modification of these parameters required: (a) identifying the corresponding atom names from the 3D structure file, (b) mapping the atom names to atom types stored in topology file, (c) finding the corresponding information in the parameter file by using the atom types, and (d) applying the modifications. To investigate the configuration of deoxycumambrin, six improper dihedral angles had been disabled, and two new improper angles were added by repeating the manual step, ‘*Topology/parameter adjustments*’. The force constants associated with the bonds, angles, and proper and improper dihedral angles needed to be adjusted in the parameter file to match the Xplor-NIH calibration of relative energy terms, and this was accomplished in the manual step, ‘*Adjusting energy parameters*’. In addition, by default, Xplor-NIH requires the atoms in PDB ligand files to be identified with an ‘ATOM’ tag, whereas most programs, including PRODRG, report the atoms with ‘HETATM’ tags. Moreover, Xplor-NIH requires atom names to be left-aligned in the input PDB file, this was accomplished by the manual step, ‘*Adjusting PDB file format*’. The final manual step of the workflow was to convert the RDC restraints from the standard format to the Xplor-NIH input format (‘*Preparing restraint*s’).

Figure 5b shows the structural investigation of deoxycumambrin as carried out under the RUNER framework. We used the same input structure file and uploaded the RDC restraints to the RUNER web server. From the types (Amber, CHARMM, Xplor-NIH) of output returned to the RUNER website, we chose those items necessary for molecular modeling with Xplor-NIH: (a) a PDB file formatted according to Xplor-NIH expectations, (b) an RDC file in a well-defined format, and (c) force field parameters stored in a pair of topology and parameter files. We note that the atom naming system was consistent in these files, therefore no manual adjustment was needed. The modification of the force field parameters was conducted using the interactive interface of the RUNER GUI. There was no need to identify the corresponding atom types or search through the topology and parameter files, because, as mentioned above, users readily visualize the atom names associated with the 3D structure of the compound and can apply modifications directly on the GUI.

**Fibrosterol sulfate A**. The investigation of the fibrosterol linker configuration followed the same manual steps (Figure 5a), except that a set of NOESY restraints were utilized together with a set of^1^D_CH_ RDC restraints.^1^ During the ‘*Atom label mapping*’ step, labels of 84 hydrogen atoms of fibrosterol experimental restraints were manually mapped to PRODRG output structure. Because the atom labels of heavy atoms of the input structure were not compatible with the Xplor-NIH format, atom labels of both the heavy atoms and hydrogen atoms were converted to the acceptable format during the ‘*Atom label conversion*’ step. The tedious manual parameter adjustment was performed on the relatively large natural product with 161 atoms. During this step, three improper dihedrals had to be disabled and one proper dihedral angle changed to an improper angle. The last three manual steps (‘*Adjusting energy parameters*’, ‘*Adjusting PDB file format*’, ‘*Preparing restraints*’) were performed as discussed for deoxycumambrin.

The procedure for structure calculation with the RUNER pipeline was the same as with deoxycumambrin, except that both NOE and RDC files were uploaded to the RUNER web server to keep the atom names consistent and to convert their formats to Xplor-NIH. The above two examples demonstrate how the RUNER pipeline can reduce days of error-prone manual effort to a matter of hours.

## Discussion

Drug design and screening studies frequently involve molecular modeling of natural products, drug-like fragments, and metabolites. We have identified several challenges associated with the preparation of input files for commonly used molecular modeling software packages such as Xplor-NIH. We have developed a pipeline that overcomes these challenges by utilizing the unique atom labeling program (ALATIS) as the input to the antechamber and MOPAC molecular mechanics packages. The outputs from this pipeline contain the atom specific topology and force field parameters necessary for performing molecular modeling on the compound by Amber, CHARMM, and Xplor-NIH. This module of the RUNER pipeline is publicly available as a web server that takes advantage of the high-throughput computing platform HTCondor to facilitate multiple submissions. We have demonstrated applications of this module on more than 4,100 small molecules that include a set of FDA approved drugs and a set of natural products. As part of the RUNER pipeline, we have introduced a standalone graphical user interface that provides seamless modification of input topology and parameter files for Xplor-NIH. We illustrate the functional features of the user interface through its application to structural studies of two natural product compounds. The data set for force field parameters of small molecules complements a growing database of standardized small molecule data available from our website [http://runer.nmrfam.wisc.edu/].

### Availability of the code

The RUNER web server is publicly available to academic users for performing the computational portion of the pipeline. The RUNER GUI for modifying input files to Xplor-NIH is available through the NMRBox^36^ project [https://nmrbox.org/], which is freely available to academic users. In addition, the GUI, which was developed using MATLAB®, is available on the RUNER website [http://runer.nmrfam.wisc.edu/]. This work is copyrighted under the terms of GPL. The web-service and the GUI are provided on an ‘as is’ basis without warranty of any kind, either expressed or implied. All academic uses of the web server, or modification and application of RUNER, are free provided that the corresponding publications are cited.

## Acknowledgment

This study made use of the National Magnetic Resonance Facility at Madison, which is supported by National Institutes of Health (NIH) grant P41GM103399 (NIGMS) and the BioMagResBank (BMRB), which is supported by NIH grant R01GM109046. H.D., J.R.W., and H.R.E. were supported in part by the National Center for Biomolecular NMR Data Processing and Analysis, which is supported by NIH grant P41GM111135 (NIGMS). CDS is supported by the Intramural Research Program of the Center for Information Technology at the NIH.

